# Social foraging in vampire bats is predicted by long-term cooperative relationships

**DOI:** 10.1101/2021.04.23.441116

**Authors:** Simon P. Ripperger, Gerald G. Carter

## Abstract

Stable social bonds in group-living animals can provide greater access to food. A striking example is that female vampire bats often regurgitate blood to socially bonded kin and nonkin that failed in their nightly hunt. Food-sharing relationships form via preferred associations and social grooming within roosts. However, it remains unclear whether these cooperative relationships extend beyond the roost. To evaluate if long-term cooperative relationships in vampire bats play a role in foraging, we tested if foraging encounters measured by proximity sensors could be explained by wild roosting proximity, kinship, or rates of co-feeding, social grooming, and food sharing during 22 months in captivity. We assessed evidence for six hypothetical scenarios of social foraging, ranging from individual to collective hunting. We found that female vampire bats departed their roost individually, but often re-united far outside the roost. Nonrandomly repeating foraging encounters were predicted by within-roost association and histories of cooperation in captivity, even when controlling for kinship. Foraging bats demonstrated both affiliative and competitive interactions and a previously undescribed call type. We suggest that social foraging could have implications for social evolution if ‘local’ cooperation within the roost and ‘global’ competition outside the roost enhances fitness interdependence between frequent roostmates.

## Introduction

Socializing and foraging are two key determinants of reproduction and survival that can influence each other in several interesting ways. Preferred social relationships can drive foraging decisions (e.g. great tits: Firth et al. (2015)). Conversely, shared foraging behaviors might shape how relationships form (e.g. bottlenose dolphins: Machado et al. (2019)). Social relationships can determine access to food because closely affiliated individuals can peacefully co-feed at a food patch, hunt together (Lang and Farine, 2017), cooperatively defend food patches (e.g. Emery et al., 2007; Robichaud et al., 1996; Seed et al., 2008), or even give food to less successful foragers (e.g. chimpanzees: Samuni et al. (2018)). Access to food is therefore one benefit of long-term *cooperative relationships*, i.e. stable preferred associations that involve cooperative investments such as grooming and food sharing. For example, grooming in chacma baboons promotes tolerance during foraging (King et al., 2011), and vervet monkeys strategically groom individuals that control access to food due to social dominance (Borgeaud and Bshary, 2015) or an experimentally manipulated ability to access food (Fruteau et al., 2009). A particularly clear non-primate example of cooperative relationships providing food occurs in common vampire bats where females regurgitate ingested blood to socially bonded kin and nonkin that failed to feed that night (Carter and Wilkinson, 2013; Wilkinson, 1984).

Food-sharing relationships in vampire bats form as preferred associates escalate social grooming (Carter et al., 2020). These preferred associations and cooperative interactions occur within the day roost. However, little is known about if or how cooperative relationships extend beyond the roost. For example, foraging with socially bonded roostmates might increase efficiency in searching for prey or feeding from wounds, but it remains unclear if or how vampire bats perform social hunting. Several authors provide anecdotal evidence for: groups of females apparently flying together, adult females departing roosts in groups of 2-6, and groups arriving together at a pasture, or approaching and circling prey (Crespo et al., 1974; Greenhall et al., 1971; Wilkinson, 1985; Wilkinson, 1988). There are also observations of up to four individuals feeding simultaneously from different wounds on the same cow (Greenhall et al., 1971), or pairs feeding on the same wound (Greenhall et al., 1971; Wilkinson, 1985). Wilkinson (1985) described evidence that mother-daughter pairs co-forage and share wounds, but found no evidence that frequent roostmates forage together.

Social foraging can take many forms, from mere aggregations attracted to a common resource to coordinated foraging groups with differentiated roles. Socially-hunting species can be placed on a spectrum of resource sharing from individual foragers competing to group-level sharing (Lang and Farine, 2017). The form of social foraging and the scale of competition over resources outside the roost can have implications for the evolution of food-sharing relationships. Several evolutionary models of vampire bat food sharing as multi-level selection view them as foraging individually then sharing food at the group-level (Di Tosto et al., 2007; Foster, 2004; Witkowski, 2007), but this view contrasts with evidence that food-sharing relationships within groups are reciprocal and highly differentiated (Carter and Wilkinson, 2015; Wilkinson, 1984). An alternative possibility is that individualized relationships drive both within-roost resource sharing and social hunting. This hypothesis is not mutually exclusive with group hunting, because even if individuals forage in groups, specific pairs could be more likely to compete or share a wound or host (Delpietro et al., 2017; Greenhall et al., 1971; Wilkinson, 1985; Wilkinson, 1988).

Here, we assessed the relative evidence for a range of hypothetical scenarios that vary in degree of coordination of social foraging among socially bonded bats (Figure 1). Preferred roostmates might not coordinate their behavior outside the roost. If instead bats optimize individual foraging efficiency by preferentially depart, follow, or forage with their preferred roostmates, then within-roost networks should predict co-departures or foraging encounters. Alternatively, to maximize their collective search area, bats might prefer to forage with bats outside their network of cooperative relationships and actually avoid foraging with their frequent roostmates. If so, within-roost and outside-roost networks should be negatively correlated. Finally, if entire roosting groups also forage together, then we expect highly correlated within-roost and outside-roost networks.

**Figure 1.**
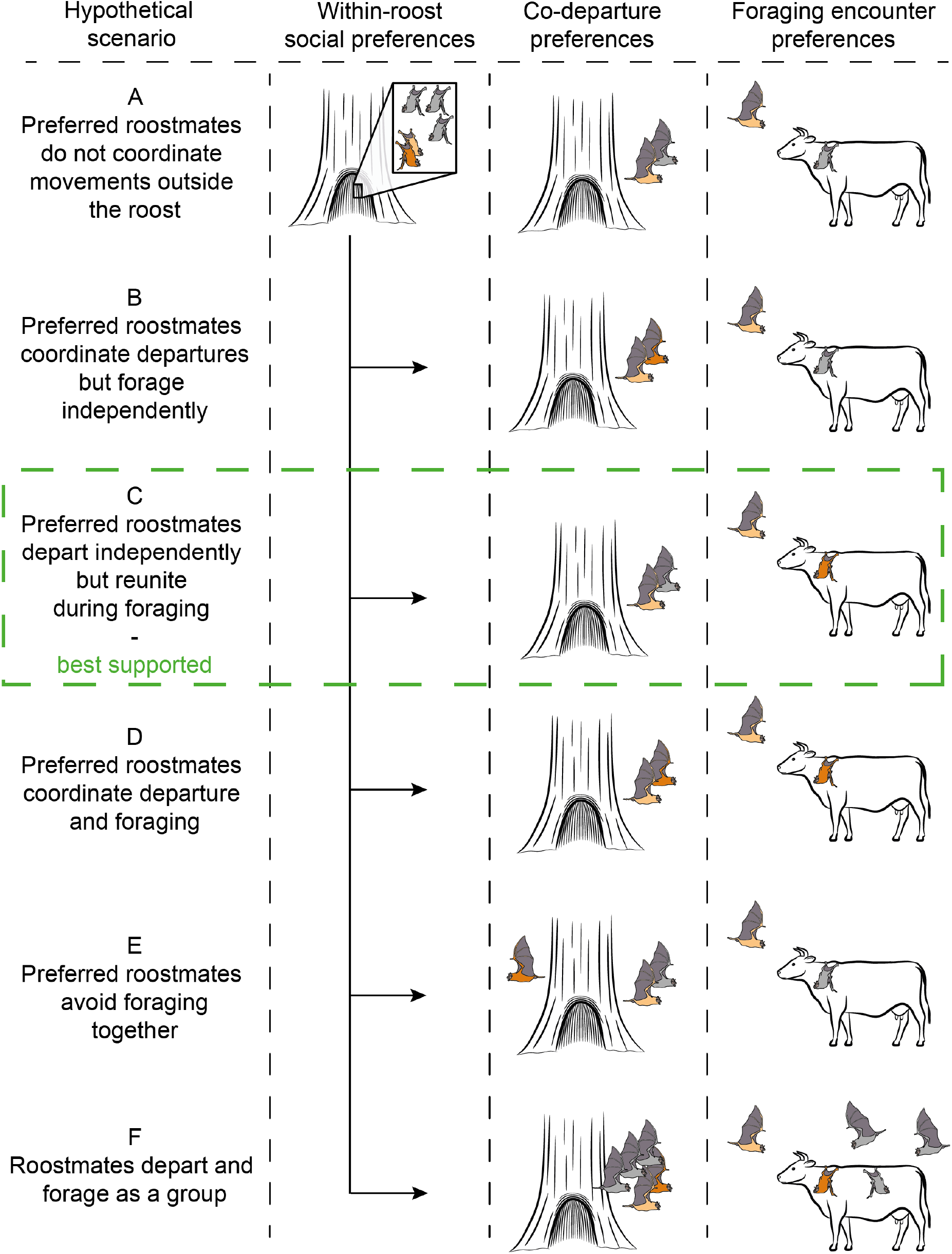
Hypothetical scenarios for how within-roost relationships predict foraging. For the same roosting association networks, each scenario predicts different outcomes for how preferred roosting relationships correlate with co-departures or encounters during foraging. Preferred roostmates (shown as pair of light brown and dark brown bats) might either: not coordinate their behavior outside the roost (A), coordinate only their departures (B), depart independently and then reunite during foraging (C), coordinate departures and foraging (D), or avoid foraging together (E). Alternatively, the bats could depart and forage as a large group (F).

To evaluate evidence for these scenarios, we tested whether nightly foraging departures and encounters were predicted by: kinship, roosting associations based on two levels of proximity (during the previous day or over the whole study), and rates of social grooming, food sharing, and co-feeding in captivity. To document roosting associations and foraging encounters, we analyzed social encounter data from proximity sensors placed on 50 free-ranging vampire bats. As additional predictors for 23 of these bats, we used unpublished data on captive co-feeding rates and published long-term rates of social grooming and food sharing (Ripperger et al., 2019). Using simultaneous ultrasonic recording and infrared video, we also describe a distinct new type of vampire bat call only observed during hunting interactions. Our findings illustrate how within-roost cooperative relationships influence foraging in vampire bats and how social networks can vary across contexts.

## Methods

### Subjects

Subjects were common vampire bats (*Desmodus rotundus*) including 27 wild-caught adult females that were tagged and released, and 23 previously captive females (17 adults and their six subadult captive-born daughters) that had spent the past 22 months in captivity and were then tagged and released back into their wild roost tree (see Carter et al., 2020; Ripperger et al., 2019). See supplement for details.

### Kinship

We assumed that known mother-daughter pairs had a kinship of 0.5. To estimate kinship for all other pairs, we genotyped bats at 17 polymorphic microsatellite loci (DNA isolated via a salt–chloroform procedure from 3-4 mm biopsy punch stored in 80 or 95% ethanol), then used the Wang estimator in the R package ‘related’. See supplement for details.

### Past cooperative interaction rates in previously captive bats

To measure cooperative relationships in the previously captive bats, we used previously published rates of social grooming and food sharing from experimental fasting trials (Carter et al., 2020). See supplement for details. To assess tolerance while feeding, we also analysed previously unpublished data on co-feeding among the same captive vampire bats. Social interactions were observed at blood spout feeders while the bats were in captivity, including 1300 competitive interactions and 277 cases of co-feeding where two bats were observed feeding from the same blood spout at the same time (from 1050 h of observation from 70 nights). We used 201 co-feeding events with identified bats to construct a co-feeding network of the number of dyadic co-feeding events (range = 0 to 6) for each pair.

To assess correlations between the captive co-feeding network and networks of food sharing or social grooming, we used Mantel tests. To test the same correlation while controlling for overlap in individual feeding times, we also used a custom double permutation test (Farine and Carter, 2020). This procedure calculates an adjusted co-feeding rate for each pair as the difference between the observed co-feeding rate and the median expected co-feeding rate from 5000 permutations of the co-feeding bat identities, permuted among the bats seen within each hour. The results of this constrained permutation test and the unconstrained Mantel test were similar and gave the same conclusion, so we report only the results from the double permutation test. To test for preferred captive co-feeding partners, we also used the same within-hour permutations to test if social differentiation in co-feeding (the coefficient of variation in co-feeding rates) was greater than expected from the null model.

### Association rates in the wild using proximity sensors

We placed custom-made proximity sensors on all 50 female common vampire bats (sensor mass: 1.8 g; 4.5-6.9 % of each bat’s mass) that automatically documented dyadic associations among all 50 tagged bats when those come within reception range (max. 5-10 m). To log encounters, each proximity sensor broadcasted a signal every two seconds to update the duration of each encounter. We used 1 s as the duration of encounters that were shorter than two successive signals (i.e., encounters shorter than two seconds). The maximum signal strength of each encounter can be used as an estimate for a minimum proximity between two tagged bats during the encounter by comparing the signal intensity to a calibration curve (Ripperger et al., 2019; Ripperger et al., 2020b).

We collected association data on the free-ranging bats at Tolé, Panama (8°12’03”N 81°43’46”W), a rural area that is mainly composed of cattle pastures for meat production. Around 200-250 common vampire bats roosted inside a hollow tree on a cattle pasture that was about 15 ha in size. To create a stable food patch, we corralled ca. 100 heads of cattle at a distance of ca. 300 m from the roost from 6pm until 6am between the evening of September 21 until the morning of September 26, 2017 (days 1 to 5 in our study). Before and after that time period, the cattle were ranging freely. A neighboring, much larger pasture west of the roost had about 1,500 heads of cattle within a distance of 1-2 km (Figure S1).

To construct networks of roosting association rates during each daytime period within the roost, we relied on roosting association data that had been used in a previous study (Ripperger et al., 2019). Based on the same two thresholds of signal strength as before, we defined two categories of proximity: “associations” (within a maximum of ca. 50 cm) and “close contacts” (within ca. 2 cm). Roosting network edges were rates of within-roost association or close contact, i.e. the total time two bats spent in association per unit of time. See supplement for details.

To log presence and co-occurrence of foraging bats at points outside the roost during the night, we placed base stations (which can detect tagged bats at distances of about 150 m) at the roost and at 5 other locations in the surrounding cattle pastures. To identify departures from the roost, we found the points in time where each bat lost connection from the roost base station and almost all of the many tagged bats in the colony within communication range (i.e., a sudden drop in associations from many bats down to 0-3 bats; see figure 2 in Ripperger et al. (2020b)). Departing bats may have also contacted base stations on the cattle pasture (Figure S1). We used the same kind of data to find the return times to the roost for each bat and night.

**Figure 2:**
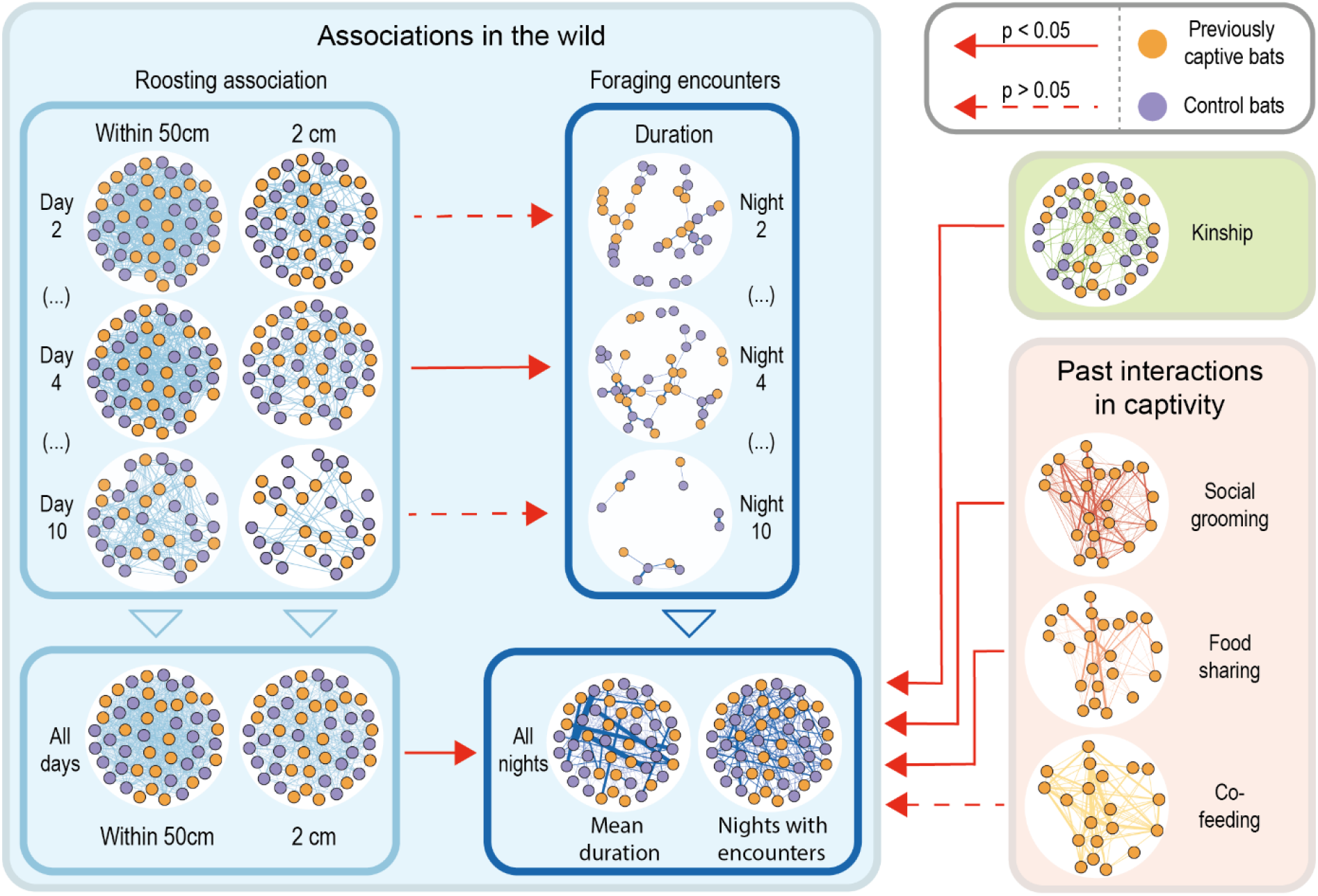
Network comparisons. Foraging encounter rates were predicted by roosting associations, kinship, and previous long-term rates of social grooming and food sharing in captivity. To facilitate visual comparisons, we fixed the spatial coordinates of each node (except for the sparse night-by-night foraging networks), we scaled edge strength in each network, and we removed nodes without edges. In the kinship network, only edges with kinship estimates > 0.24 are shown and bats without kin in the group are not shown. In the paired night-day networks of association in the wild, we only detected a clear correlation between day and night on day 4 (Table S1).

Of the 629 dyadic encounters that occurred one minute after leaving the roost and one minute before arriving at the roost, we excluded 43 encounters from further analysis, because a proximity sensor contacted the roost base station, suggesting that those encounters occurred while bats were roosting at the entrance or on the outside of the roost tree. The remaining 586 encounters occurred farther away, outside the communication range of the roost base station, and we refer to these as “foraging encounters”.

### Observing interactions of foraging vampire bats

At Tolé, we only observed two occasions, where two bats stopped at the cattle pasture and were associated (for 3.5 and 4.6 minutes). When releasing the corralled cattle in the morning we observed bite marks but to avoid changing their behavior, we did not get close enough to the cattle at night to record audio or video of bats interacting. To collect direct observations on foraging behavior, we recorded simultaneous audio and video of bat foraging behavior at a different farm near La Chorrera, Panama (8°52’42”N 79°52’05”W) using an infrared (IR) spotlight, IR-sensitive video camera (Sony AX53 4K camcorder) and a Avisoft condenser microphone (CM16, frequency range 1 to 200 kHz) and digitizer (Avisoft USG 116Hbm, 1000 kHz sampling rate, 16-bit resolution) connected to a notebook computer. One observer (SPR) moved with a herd of about 20 grazing cattle without visual light and observed the moving cattle through the viewfinder of the IR camera. To compare social calls made during foraging with calls from inside a roost, we used the same recording equipment to record social calls from a roost only a few hundred meters from the foraging site.

### Acoustic analysis of calls in foraging bats

We used Avisoft SASLab Pro (R. Specht, Avisoft Bioacoustics, Glienicke/Nordbahn, Germany; version 5.2.13) to measure acoustic parameters of the social call types. Start and end of calls were determined manually, based on the oscillogram. Subsequent, five acoustic parameters were measured automatically; one temporal (duration) and four spectral parameters (peak frequency at maximum amplitude, minimum and maximum frequency, and bandwidth). Acoustic parameter extraction was restricted to the fundamental frequency.

Spectrograms were created using a Hamming window with 1024-point fast Fourier transform and 93.75 % overlap (resulting in a 977 Hz frequency resolution and a time resolution of 0.064 ms). To estimate the frequency curvatures of the different call types, we measured the spectral parameters at 11 different locations distributed evenly over the fundamental frequency of each call. To compare call structure from different contexts (roosting vs foraging, antagonistic vs affiliative behavior) in multivariate space, we plotted the first two principal components after entering these measures into a principal component analyses with varimax rotation (using the ‘foreign’ package in R).

### Statistical analysis of foraging behavior

For every bat, we calculated the times over 9 days when it was clearly distant from the roost tree (ESM File 2). To test whether the previously captive bats and never-captive control bats differed in the departure time and duration of their foraging bouts, we fit linear mixed-effect models (LMMs) with type of bats and day as fixed effects and bat as a random effect (p-values estimated with Satterthwaite’s degrees of freedom method using the R package lmerTest). To compare consistency of onsets and durations, we measure the unadjusted repeatability (ICC or intra-class correlation) for each type of bat. To see how often tagged bats departed together, we inspected cases where departure times were within one minute.

### Preferred associations during foraging

To test if repeated foraging encounters occurred among the same bats more than expected by chance, we used a custom data permutation test that compared observed and expected social differentiation (the coefficient of variation in co-foraging rates, which increases when some pairs have more repeated encounters) while controlling for overlap in foraging times. Since all bats were sampled evenly within each night and most foraging encounters were brief (median = 1 second), we first used a simple and conservative measure of co-foraging rates based on counting the presence or absence of an encounter during each hour outside the roost over 9 days (counts varied from 0 to 15). For instance, if two bats met twice in the same hour bin, this is still one encounter. These binary observations could be swapped in our null model. To generate a null distribution of 5000 social differentiations expected by chance, we permuted one bat in every dyad to a random possible partner that was also present outside the roost in that same day and hour (to control for overlaps in foraging times).

### Predictors of social foraging

To test predictors of social foraging, we constructed foraging encounter networks where edges were based on either duration of total encounter time outside the roost (seconds) or number of days with foraging encounters (0-9). The latter response variable is far more conservative because it only counts repeats across different days. We included the following predictors: kinship, two proximity levels of within-roost association, social-grooming rate, food-sharing rate. We also tested the effect of dyad type (i.e. both bats previously captive, both bats never captive, one bat previously captive, and both bats captive-born juveniles). We did not use number of nights with foraging encounters as a response for tests that only included the previously captive bats, because 9 of them (including all captive-born bats) left the roost during the study period (Ripperger et al., 2019). To measure how much longer foraging encounters were between kin versus nonkin, we fit a linear mixed effects model with log-transformed duration as the response variable, kinship greater than 0.1 as a binary fixed effect, and both bats’ identities as random effects, then converted model coefficients into a percentage difference.

To test the effect of predictor networks on a response network, we used regression quadratic assignment procedure (QAP) for single predictors, or multiple regression quadratic assignment procedure with double semi-partialling (MRQAP) for two predictors (using the ‘asnipe’ R package (Farine, 2013)). To create null models, we used constrained (within-day) node-label permutations. This approach is necessary for preserving the daily and nightly network structure (e.g. distribution of edges and edge weights) and for controlling for the presence or absence of bats in the roost each day. To control for foraging bout overlap in each pair, we included that measure as a covariate. We also used QAP to test whether the within-roosting association on each day, predicted the subsequent foraging network that night. We then bootstrapped the mean of the slopes across the eight days to test for an overall paired day-night effect.

### Consistency of individual social traits

To test whether bats that are more socially connected within the roost are also more connected in foraging networks, we tested if the nodes’ degree centrality was correlated between roosting and foraging networks. We measured degree centrality independently within each day or night network when the bat was present and then took the mean for each bat. Bats with no encounters in that day or night were considered missing for that day (i.e. not counted as zero degree). We fit general linear mixed effect models with foraging network centrality as the response variable, roosting network centrality (either association and close contact) as fixed effect, and bat as random effect. P-values were calculated from 5000 permutations of the bat’s foraging centralities within each night. These constrained node-label permutations (within night) are necessary to control for the fact that foraging and roosting network centralities could be correlated simply by some bats being present at the site longer.

## Results

### Sampled bats did not depart together

The never-captive control bats departed from the roost 8.3 hours after sunset and returned 2.5 h later, on average (ESM 2). The previously captive bats foraged earlier and less predictably (see below). We observed only five cases where two bats departed within five seconds of each other and none of these cases was followed by a foraging encounter. For the cases where pairs did have a foraging encounter, the shortest differences in departure times were 8, 21, and 28 s.

### Previous captivity influenced departures and foraging

Compared to the wild control bats, the previously captive bats departed the roost on average 1.6 hours earlier (t = -4.55, p<0.0001), but they did not forage consistently longer (LMM; t=1.29, p=0.2, ESM 2). The captive-born bats departed 2 hours earlier (t = -3.15, p = 0.002) and also did not forage longer (t = -0.41, df = 47.8, p = 0.7) than control bats. All these models control for departure times being on average 14 minutes later each day (t = 6.6, p < 0.0001, ESM 2), perhaps due to moonset times being 20-40 min later each day during the study period. The total duration of foraging encounters did not clearly differ between types of pairs (Figure S3A), but pairs of control bats had significantly more nights with foraging encounters (Figure S3B) compared to other types of pairs, possibly due to control bats having more consistent foraging times. Departure times were more consistent across days within each control bat (intraclass correlation coefficient (ICC) = 0.58) compared to within each previously captive bat (ICC = 0.21) or captive-born bat (ICC = 0). The duration of the longest foraging bout was also more consistent in wild control bats (ICC = 0.54) than the previously captive bats (ICC = 0.35) or captive-born bats (ICC = 0.15).

### Preferred associations in foraging encounter networks

Foraging encounters were orders of magnitude shorter in duration than within-roost encounters, their median duration was 1 s, and they never exceeded 30 minutes (ESM 1, Figure S2). Of 151 pairs with a foraging encounter, 45 did this repeatedly across 9 nights. The variation in number of hours in which two bats reunited was greater than expected from our null model that simulated random encounters among bats that were outside the roost in the same hour (observed coefficient of variation = 4.36; p < 0.001; 95% of expected values: -2.2 to 2.4). Most of these foraging encounters occurred at locations outside our sampled areas, but 10 (among eight pairs) occurred near the other base stations on the surrounding cattle pastures (ESM 1, Figure S1), and only three foraging encounters (among three pairs) occurred at the corral that we created as a stable food patch about 300 m from the roost (two encounters on days one and three while the cattle were present and one encounter on day seven).

### Kinship predicts foraging encounters

Kinship predicted the number of nights with foraging encounters (QAP, β = 15.4, n = 46 bats, p < 0.0001) and foraging encounter time (β = 15.4, n = 47 bats, p = 0.022) even when controlling for bout overlap (MRQAP, β = 0.10, p = 0.002). The median duration of a foraging encounter for close kin (r > 0.1) was 9 s, which was 135% longer in duration relative to the duration of foraging encounters between nonkin (r < 0.1; median duration = 1s; β =0.85, df=175, p=0.001).

### Within-roost association rates predicted foraging encounters

Bats that spent more time near each other within the tree during the day, also spent more time together outside the roost during the night (associations: QAP, β = 29.5, p < 0.001; close-contact: QAP, β = 24.7, p = 0.002) even when controlling for the foraging bout overlap (associations: MRQAP, β = 0.092, p = 0.003; close-contact associations: MRQAP, β = 0.078, p = 0.015). Within-roost associations also predicted a greater number of nights with foraging encounters (associations: QAP, β = 0.07, p < 0.001; close-contact association: QAP, β = 0.04, p = 0.021). The relationship between the within-roost association network and the corresponding night’s foraging network was clear within only one of the 8 days (Table S1), but the paired relationships between day and night networks tended to be greater than zero overall (associations: mean β = 0.026, 95% CI = 0.004 to 0.051; close-contact associations: mean β = 0.018, 95% CI = 0.003 to 0.04).

### Roosting degree centrality predicted foraging degree centrality

Bats that were connected to more associates in within-roost association networks also tended to have more associates in the nightly foraging networks (associations: β = 0.034, n = 48 bats, one-tailed p = 0.008; close-contact: β = 0.055, one-tailed p = 0.078; Figure S4).

### Cooperative relationships in captivity predict foraging encounters in the field

In the previously captive bats, foraging encounter time was predicted by social grooming (QAP, β = 26.5, n = 22, p = 0.032; MRQAP controlling for bout overlap: β = 0.12, p = 0.063) and food sharing (β = 38.7, n = 22, p = 0.015; MRQAP controlling for bout overlap: β = 0.20, p = 0.014), and food sharing when controlling for kinship (MRQAP, sharing: β = 0.16, p = 0.022; kinship: β = 0.14, p = 0.049). Kinship and cooperative relationship were therefore independent predictors of social foraging.

### Co-feeding among familiar captive bats was not limited to cooperative relationships

In contrast to the measures of social foraging in the field, we detected only weak evidence for preferred associations during co-feeding in captivity (social differentiation = 2.10, p = 0.047 when controlling for hour, p = 0.041 when not controlling for hour), and we found no correlation between captive co-feeding and social grooming (r=0.008, p=0.36), food sharing (r=0.015, p=0.28) or social foraging time in the wild (r=0.003, n=20, p=0.42).

### Behavioral interactions during foraging

To record a sample of bat interactions during foraging encounters, we recorded infrared video and ultrasonic audio of 14 interactions between foraging vampire bats (Tables S2 and S3). Social calls during foraging had three general spectral shapes (Figures S5 and S6): “downward sweeping calls” have been recorded often in roosts and are produced by socially isolated vampire bats in captivity (Carter et al., 2012; Carter and Wilkinson, 2016). “Buzz calls” were noisy without clear tonal structure and occurred during antagonistic interactions. We observed “n-shaped calls” produced by bats interacting while near cattle (Figure S6), and to our knowledge this call type has never been seen in wild roosts, from confrontations at the feeders in captivity (Sailler and Schmidt, 1978), or from individually isolated bats in captivity (Carter et al., 2012, Carter, unpublished data).

## Discussion

### Long-term cooperative relationships predicted repeated foraging encounters

Tagged female vampire bats departed the roost individually, but often re-united far from the roost during foraging bouts. The rates of these foraging encounters were consistently higher than expected in specific pairs and predicted by roosting associations, kinship, and by the history of social grooming and food sharing in captivity, even when controlling kinship. Previous experiments with female vampire bats suggest that these measures—roosting proximity, social grooming, and food sharing—reflect an underlying cooperative relationship (Carter et al., 2020; Carter and Wilkinson, 2013; Ripperger et al., 2019; Wilkinson, 1984; Wilkinson, 1985). In this study, we knew the cooperation histories among the previously captive bats, and that these individuals had no interactions with the control bats for at least the previous 22 months. We could therefore infer that relationships that are typically defined by associations and cooperative interactions within roosts, also extend beyond the roost and may provide benefits during foraging. In addition to consistent social relationships across context (from captivity to roosting to foraging), we found that bats that encountered more associates in the roost during the days also encountered more associates while foraging during the nights, suggesting consistent individual variation in social traits.

Although some foraging encounters may have occurred before or after foraging, most of these encounters were likely to have occurred during foraging for several reasons. First, foraging encounters were brief, whereas associations among non-moving bats should be much longer in duration (Figure S2). Second, foraging is likely to take up a substantial amount of the limited time outside the roost (mean = 2.4 h). After commuting, searching, and selecting a host, a vampire bat can take up to 30 minutes to select a wound site, 10-40 minutes to prepare the wound site, and 9-40 minutes to feed (Greenhall, 1972; Greenhall et al., 1971). Third, we used infrared video to observe several interactions on or near cattle that were consistent with the short durations of foraging encounters in the proximity data (e.g., Videos S1, S2, S4). Fourth, foraging encounters among close female kin had a median duration of 9 s and were longer than among non-kin (median duration of 1s), which is consistent with observations that affiliative interactions last longer.

### No clear evidence for highly coordinated collective movements

For animals with fluid social structures (e.g. high fission-fusion dynamics), it is important to clarify the ambiguous meaning of a “social group”, and similarly, one must distinguish between different possible forms of “social foraging” (Lang and Farine, 2017). In bats, the relative degree of social coordination during foraging can be difficult to assess and compare due to differing limitations in the observational methods and the lack of knowledge of differentiated social relationships within the colony. In this study, we took advantage of well described within-roost relationships to assess evidence for several alternative scenarios of foraging behavior (Figure 1). Kinship and rates of association and cooperation led to longer and more frequent foraging encounters, but we did not observe highly coordinated joint departures or collective movements (Figure 1). This fluid pattern, of not moving in coordinated stable groups yet repeatedly encountering preferred associates during foraging, is also reflected in co-roosting networks where individuals form roosting groups that frequently change composition, yet maintain preferred relationships over time (Wilkinson, 1985). Given the many unsampled bats inside the same tree (∼200), it is possible that bats departed with other unobserved roostmates, but we did not see departures of large groups (while catching bats outside the roost) nor did we see evidence for coordination between roosting and departing in the tagged bats.

The ways that specific bats reunited with preferred associates therefore remains unknown, but the downward sweeping calls that we recorded in foraging bats (Figure S6), are similar to contact calls that captive vampire bats can use to find and recognize preferred partners (Carter and Wilkinson, 2016). The role of calls, in particular a possibly foraging-specific call type (“n-shaped call” Figure S6, Figure S7), warrants further investigation. In several other bat species, there is abundant evidence for socially influenced foraging based on eavesdropping on echolocation calls (e.g. Cvikel et al., 2015; Dechmann et al., 2009; Dechmann et al., 2010; Egert-Berg et al., 2018; Lewanzik et al., 2019). Greater spear-nosed bats in Trinidad appeared to coordinate group foraging based on a group-specific contact call (Wilkinson and Boughman, 1998). Observations of the fish-eating greater bulldog bat suggested that female roostmates depart individually, then assembled into small groups outside the roost to forage together, possibly coordinating their movements with calls (Brooke 1997).

### Affiliative and competitive interactions

Given the difficulty of making a bite compared to the ease of drinking from an open wound, some individual vampire bats appear to exploit the bites already made by others, and fights can occur over open wounds or hosts (Delpietro et al., 2017; Greenhall et al., 1971; Sazima, 1978; Schmidt, 1978; Wilkinson, 1985), but it remains unclear how often these competitive interactions occur among familiar versus unfamiliar vampire bats. In our study, we observed foraging vampire bats engaging in both affiliative and competitive interactions (see Table S3, Videos S1-5). The competitive interactions were far more aggressive than what we observed among familiar captive bats feeding from an accessible and unlimited source of blood. This observation and our results above are consistent with the hypothesis that competitive interactions are more likely between less familiar bats.

### Implications for social dominance

The fluid nature of foraging encounters has potential implications for social dominance. Dominance hierarchies should be common when animals move together in groups, because the same frequent groupmates will also be primary competitors for first access to food. Dominance hierarchies among familiar female vampire bats, which do not always travel or forage together, are indeed less clear and linear than among female mammals that do travel and forage in more stable groups (Crisp et al. unpublished data). Furthermore, blood from an open wound is often not a limited resource, so competition over food might be relatively low among familiar vampire bats that tolerate each other (as observed in captivity) and even share food, compared to unfamiliar conspecifics that might “steal” a wound.

### Implications for the evolution of cooperative relationships

Vampire bats might also benefit from foraging with socially tolerant partners by acquiring information on where to feed or by gaining access to open wounds. A single open wound can sequentially feed several bats, and leading a cooperation partner to an open wound on the same host would presumably be less costly to a successful forager than regurgitating blood to that individual later at the roost. Put differently, socially bonded bats would benefit from each other’s foraging success, i.e. interdependence (Roberts, 2005). As seen in ravens, the presence of a socially bonded partner might even allow for joint defense of food against third-parties (Sierro et al., 2020; Szipl et al., 2015).

Such forms of social foraging in vampire bats may have implications for the spatial scale of competition—a key factor shaping social evolution in humans (West et al., 2006) and other group-living animals (Radford et al., 2016). In female vampire bats, cooperation occurs ‘locally’ with specific frequent roostmates, and competition over food might occur more ‘globally’ with members of the much larger population. If so, a more ‘global’ scale of competition could reduce conflict and increase interdependence among highly associated females. To test this idea, it would thus be useful to determine if sampled groups of vampire bats consistently feed on the same or different prey individuals, and if vampire bats are more likely to approach or avoid the social calls of foraging bats that are frequent roostmates versus unfamiliar conspecifics.

### Implications for describing social structure

A major advantage of proximity sensors was the ability to continuously track associations among multiple individual bats both inside and outside their roost, which allows for the construction of dynamic and multi-layer networks. Studies on social foraging and other social behaviors in bats and other small highly mobile vertebrates have historically been limited by the available tracking technology (Ripperger et al., 2020b). Radiotelemetry has poor spatial resolution and continuously tracking many individuals is difficult. Current GPS-tags for bats have rather short runtimes and the tags need to be recovered to download the data. On-board ultrasound recorders (e.g. Egert-Berg et al., 2018) do not reveal the identity of encountered individuals. A major downside to proximity sensors was that many foraging encounters occurred at unknown locations. However, placing proximity sensors or antennas at more locations and on livestock would allow a better reconstruction of foraging behavior. A combination of biologging approaches can also help to overcome existing challenges (e.g. Leoni et al., 2020; Ripperger et al., 2020a). Standardized high-throughput methods for measuring social network structure across bats and other diverse groups allow for comparative studies that assess the relative ecological and evolutionary drivers of social traits and social complexity across species.

## Supporting information

ESM1

ESM2

## Acknowledgements

Work by S. Ripperger and G. Carter is supported by a grant from the National Science Foundation (Integrative Organismal Systems #2015928). We thank R. Page and F. Mayer for providing funds for this study, which was funded by grants from the Deutsche Forschungsgemeinschaft (DFG) within the research unit FOR-1508, a Smithsonian Scholarly Studies Awards grant, and a National Geographic Society Research Grant WW-057R-17. We thank O. Castrellón and C. de León for permission to conduct fieldwork on their properties, and D. Josic, J. Berrío-Martínez, V. Flores, M. Le Chevallier, B. Cassens, N. Duda, R. Crisp, M. Nowak, and G. Cohen for their assistance during field work. We are grateful to M. Knörnschild and A. Fernandez for supporting the collection and analysis of acoustic data, I. Waurick for her valuable assistance and expertise during molecular lab work, R. Crisp for observations of co-feeding, and I. Razik and E. Siebert for creating the line drawings in Figure 1 (I.R.: cattle, tree; E.S.: bats). We thank D. Dechmann, J. Kohles, A. Fernandez, and J. Wilkinson for providing valuable feedback on earlier versions of this manuscript.

## Notes

### Competing Interest Statement

The authors have declared no competing interest.

