## Supplementary material for "Social foraging in vampire bats is predicted by long-term cooperative relationships": ESM1

### Electronic Supplementary Information 1

#### Supplementary methods

##### *Subjects: additional details*

The group of previously captive bats was captured outside the hollow tree near Tolé, Panama, in 2015 and were transferred to a flight cage at the Smithsonian Tropical Research Institute in Gamboa, Panama. Bats were marked with subcutaneous passive integrated transponders (Trovan Ltd. USA) and a visually unique combination of forearm bands (Porzana, National Tag, and birdbands.com). To feed the bats, we provided refrigerated or thawed cattle or pig blood defibrinated with sodium citrate and citric acid. To induce food sharing and grooming among the bats, we conducted 533 fasting trials over a period of 22 month. Bats were fasted 12 to 23 times (mean = 19.4 fasting trials per bat). During each fasting trial, a bat that was isolated without food for a day and night was reintroduced to the group, then observed for 1 hour. For further details see Carter et al. (2020).

The 27 wild-caught bats were mist-netted on September 19, 2017. Before sunset, we set up mist-nets in front of the roost entrance to catch exiting bats. Researchers stayed near the nets and removed bats from the mist-net immediately. Bats were kept in cotton cloth bags until we recorded sex, age and reproductive state. We fitted 27 adult females that were not visibly pregnant with proximity sensors.

##### *Kinship: additional details*

Kinship estimates were based on genotypes from 17 polymorphic microsatellite loci (markers Dr2-2, Dr6-6, Dr7-7, Dr9-9, Dr11-11, Dr15-15, Dr16-16, Dr18-18, Dr30-dr15, Dr31-dr17 (Ripperger et al., 2019, Table S2), and markers L0206, L0458, L1154, L2117, L3524, L4050, L4216 (Ripperger et al., 2021)). For the first set of markers we used a LI–COR Biosciences® DNA Analyser 4300 and the SAGA GT allele scoring software and for the second set of markers we used a SeqStudio™ Genetic Analyzer and the software Genemapper 6 to amplify and to genotype the sequences. Allele frequencies of the first marker set were based on 91 adult bats from Tolé, Panama, and 42 for the second marker set. All 17 loci passed tests for Hardy-Weinberg Equilibrium (using the R library 'genetics') and Linkage Disequilibrium (using the web version of Genepop 4.7.5; markov-chain parameters: 10,000 dememorizations, 1,000 batches, 10,000 iterations per batch). Genotypes were 99.3% complete. We chose the Wang estimator after comparing the performance of all estimators available in the package using the *simrel* function.

##### *Tracking of tagged individuals using the BATS system*

We glued custom-made proximity sensors to the bats' dorsal fur using skin-bonding latex adhesive (Perma-Type surgical bond). Tag weights were in accordance with recommendations for short-term tracking of bats (< 10 % of the body mass (Amelon et al., 2009)). We released the control group back into their roost between 4:50 am to 6:30 am on September 20th. We released the 23 previously captive bats back into the same tree at 8:12 pm.

The signal broadcasted by each proximity sensor (every 2 s) wakes every proximity sensor within 5 - 10 m from 'sleep mode', and initiates dyadic encounters between the sender and all receivers. As long as a dyad remains within reception range, the encounter duration and the maximum received signal strength indicator (RSSI) are updated every two seconds. When no signal is received from an encountered partner for 10 s (five times the sampling rate), the encounter is terminated and stored to on-board memory along with the IDs of the partners, a timestamp of the start of the encounter, encounter duration, and the maximum RSSI (as a proxy of the distance between the two partners).

##### *Identifying foraging bouts and foraging meetings*

We identified a foraging bout of a tagged bat based on the following events: (i) a sudden drop in meeting partners, (ii) an interruption of communication among the proximity sensor and the base station inside the roost, and (iii) base stations on the cattle pasture picking up the signal of the proximity sensor if the bats fly within communication range. A bat which is returning to the roost from a foraging bout should experience a sudden increase in meeting partners and establish communication to the roost base station. We therefore used the meeting architecture of every tagged bat and contact to base stations inside and outside the roost to identify clear individual foraging bouts. We used the custom-made software meeting-splitter (Ripperger, 2019), which first quantifies the number of simultaneous meeting partners for every second of the study period and then identifies situations where this number falls below a predefined number (in the present case four). We considered such events as potential foraging bouts that we further assessed visually. We confirmed potential bouts by a coinciding interrupt of the communication to the base station inside the roost that indicates absence. In addition, we checked whether the bout was flanked by or contained presence signals at base stations outside the roost confirming presence on the cattle pasture. The latter was not always true since bats may have departed from the roost without coming within communication range of base stations on the pasture and because not all base stations have been operated during all nights. This information merely presents an additional, optional layer of information. We then queried all meetings from the meeting database, which originated during the verified bouts (i.e., 'foraging encounters'). As a final quality check we verified that both participants of a foraging meeting have currently been on potential foraging bout or have at least not been identified inside the roost. Although it is possible that not all foraging bouts were identified, all foraging encounters were visually verified to have occurred outside the roost.

##### *Recording video and audio of interactions among foraging vampire bats*

On the night of June 25th, 2019, we took simultaneous and audio and video recordings of free-ranging vampire bats during foraging. The video camera, the ultrasound recorder, and the heavy-duty infrared spotlight that was powered by a 12V car battery were mounted to a tripod in a way that all three devices were facing the same direction. A Dell Latitutde E7450 PC and the car battery were carried inside a backpack. We made recordings at a distance of approximately 3-10 m and using the software Avisoft RECORDER. Gain levels were dynamically adjusted to avoid signal clipping while recording since the volume of vocalizations was highly variable depending on the distance to the recorded animals and the direction the bats were facing while emitting vocalizations. Whenever bats were spotted circling around or landing on a cow, the observer started to record video to match the ultrasonic recordings. We then used the timestamps of the synchronized audio and video to pair the two types of recordings.

In addition, we made sound recordings inside a vampire bat roost only a few hundred meters from the site where we recorded foraging bats. We used the same acoustic recording devices but sampling rate was set to 250 kHz (16-bit depth resolution). Bats were recorded at a distance of ca. 0.5-3 m and gain levels were again adjusted dynamically to avoid signal clipping.

For the acoustic analysis, we determined start and end of each call manually based on the oscillogram and used the Avisoft SASLab's automatic measurement function to extract spectrum-based parameters. Calls were multi-harmonic, and we took measurements of the fundamental frequency (first harmonic). For one call type (n-shaped calls, see results), the fundamental frequency was very faint, so we measured frequencies from the second harmonic and divided the spectrum-based measurements by 2 to obtain values for the fundamental frequency.

***Study site***

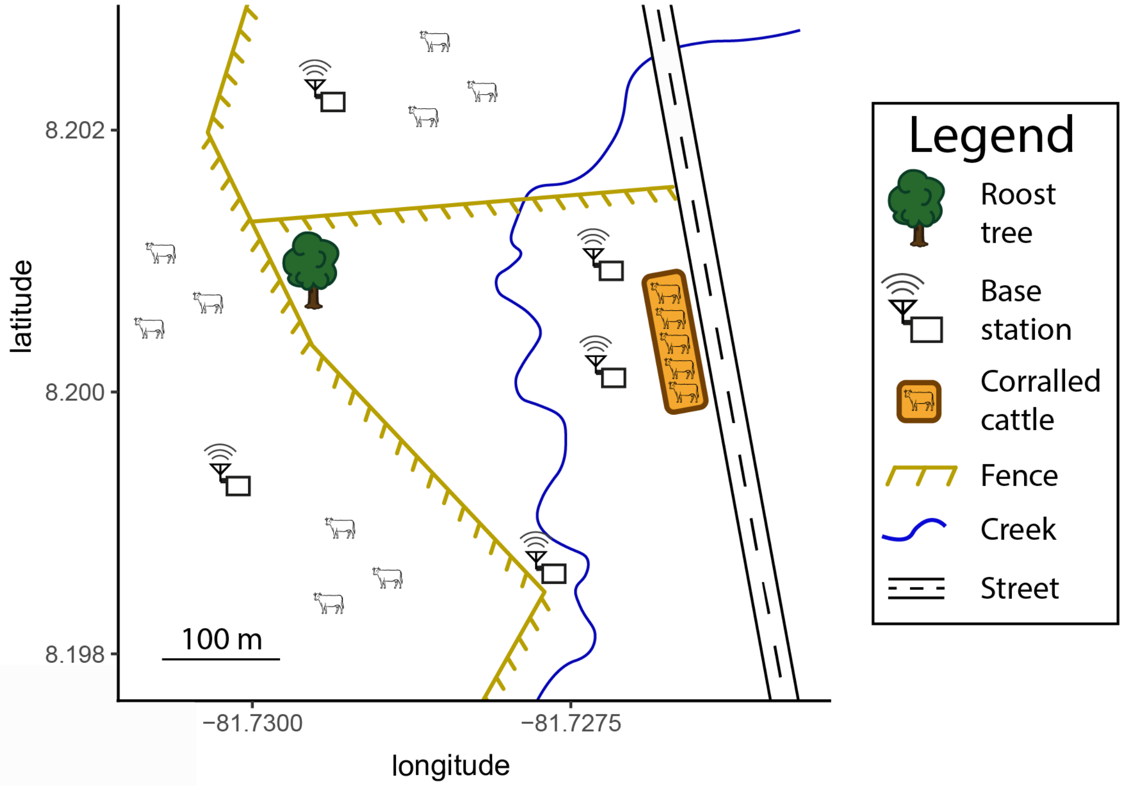


**Figure S1: Schematic map of the study site at Tolé, Panama.** Base stations could detect flying bats. Note that the corralled cattle were moving freely starting on day 6 of the study. The pastures north and west of the roost had about 1,500 heads of freely moving cattle. Line drawing of cattle by Imran Razik.

**Supplementary results (supporting figures, tables, and videos)**

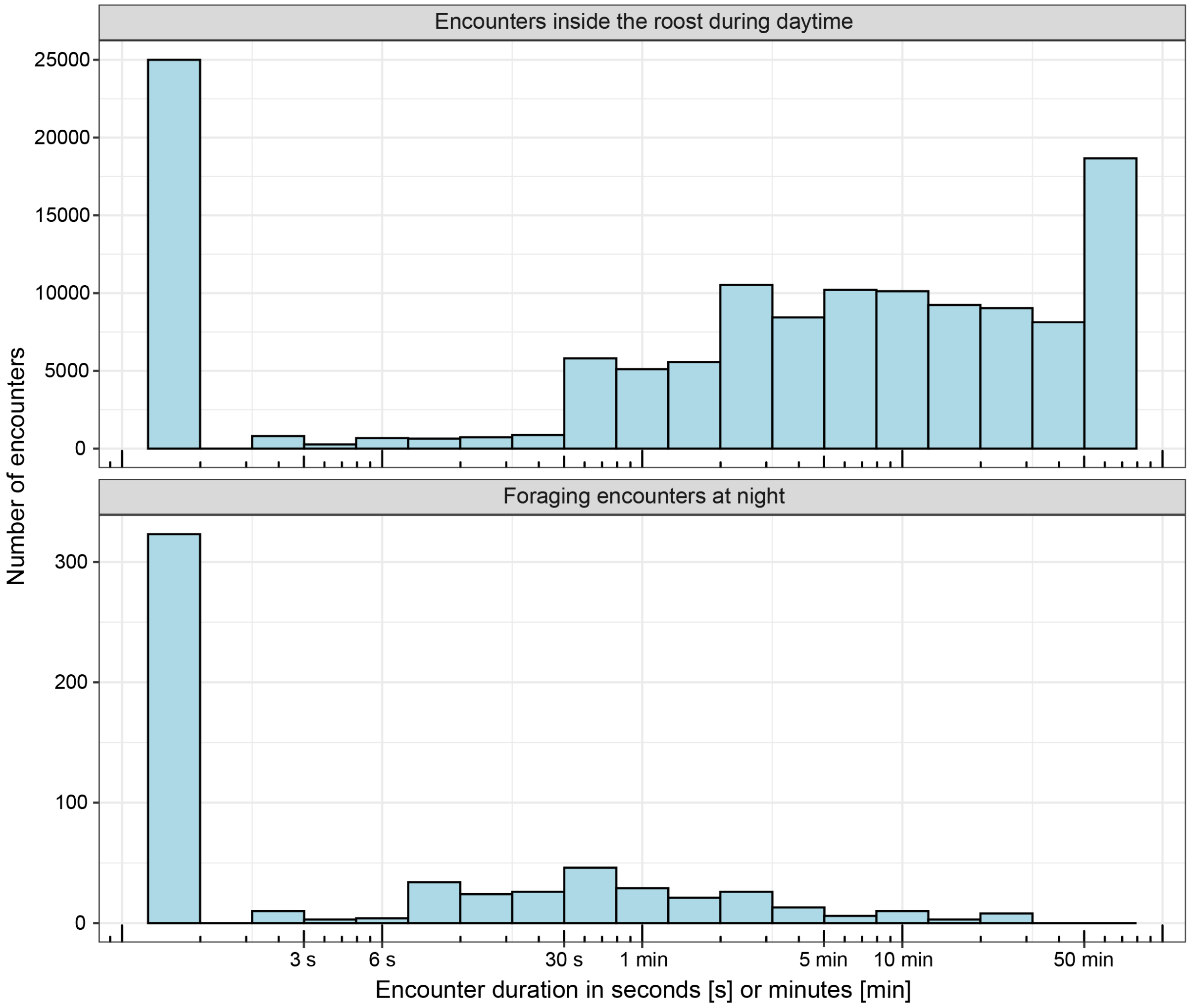


**Figure S2. Encounters inside the roost were much longer in duration than encounters outside the roost.** Encounters longer than 30 minutes were only observed inside the roost. Histograms show duration of encounters inside the roost during day (top) and foraging encounters at night (bottom). Note that time is plotted on a logarithmic scale. The high bar of short duration is one second, the shortest possible encounter duration (when bats come only briefly within sensor communication range), and the maximum duration was one hour because all encounters were divided at the hour-mark.

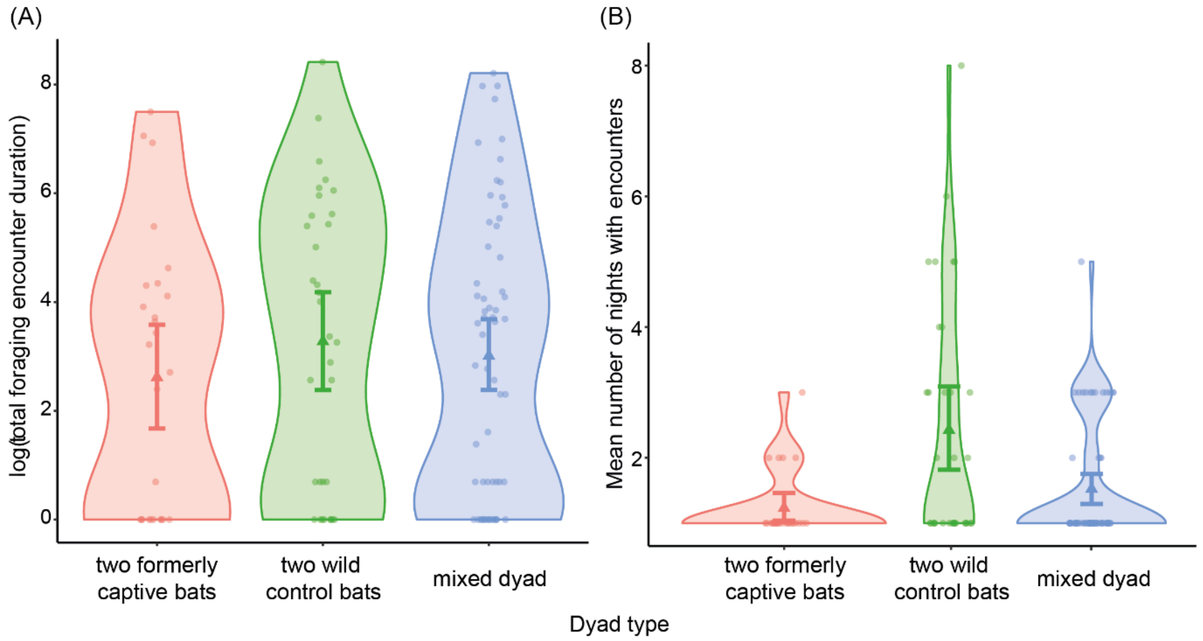


**Figure S3. Differences in social foraging by dyad type.** Violin plots show that (A) different types of dyads did not greatly differ in total foraging encounter duration (means (triangles) and bootstrapped 95% CI (error bars) for pairs of two previously captive bats = 2.61 [1.68-3.58], two wild control bats = 3.27 [2.39-4.15], and one previously captive and one wild control bat (mixed dyad) = 3.01 [2.37-3.67]), and that (B) pairs of two wild control bats had more nights with foraging encounters (two previously captive bats = 1.23 [1.07-1.42], two wild control bats = 2.42 [1.82-3.09], mixed dyads = 1.52 [1.31-1.77]).

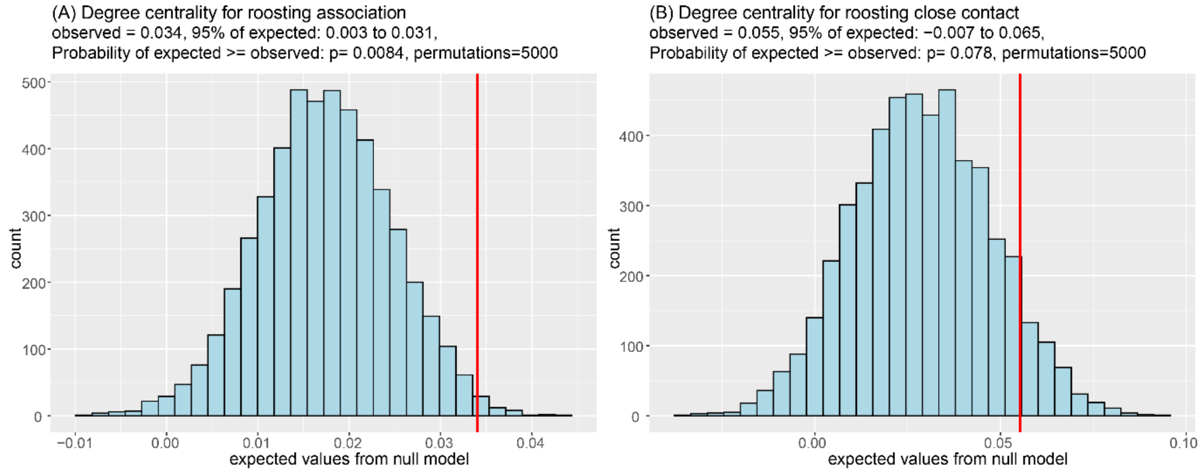


**Figure S4. Foraging degree centrality is predicted by roosting degree centrality.** Blue histograms show expected coefficients for the same model fit to randomized data generated by our null model. Red lines show the observed coefficients. Subtitles provide the observed coefficient, the 95% quantiles for the expected coefficients, the one-sided p-value, and number of permutations. Proximity of about 50 cm is required for “roosting associations” and about 2 cm proximity is needed for “close contact”, so close-contact networks are sparser.

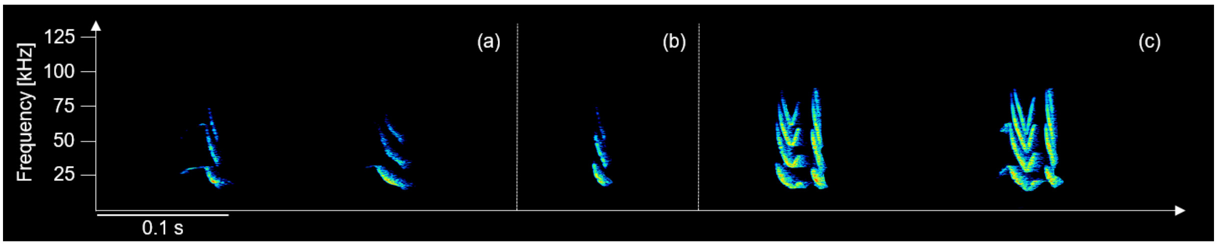


**Figure S5. Spectrograms of social calls of common vampire bats inside a roost.** Tens of bats present. Call types were (a) “undulated down sweep”, (b) “down sweep”, and (c) “u-shaped calls followed by down sweep”. Inter-call intervals and call sequences have been modified for the figure.

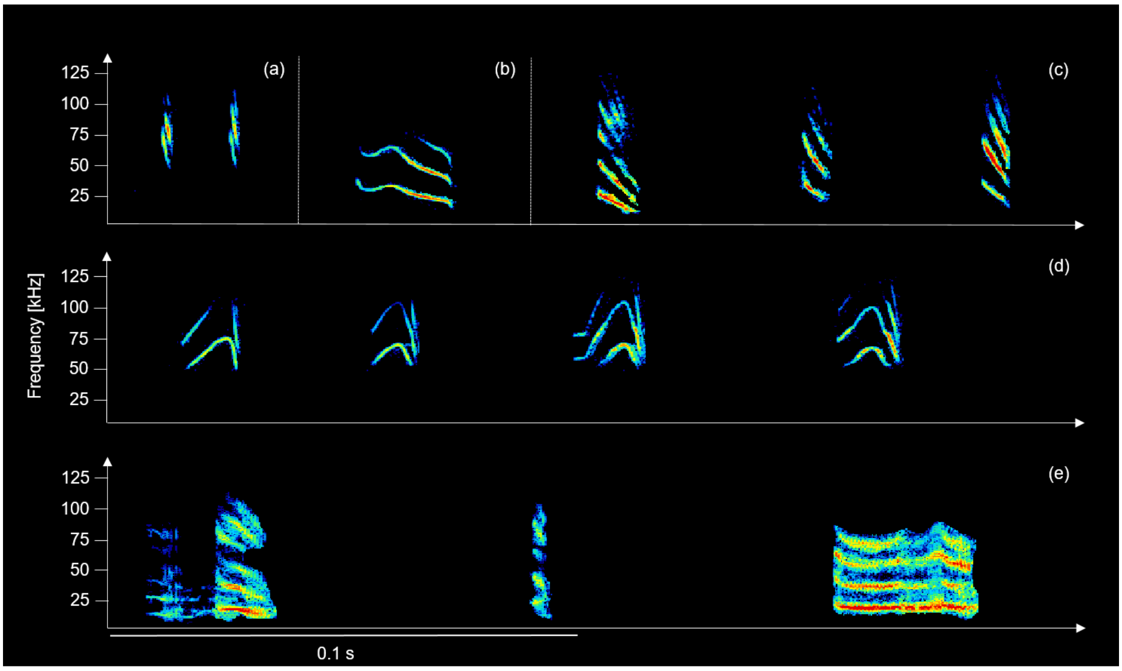


**Figure S6. Spectrograms of social calls of common vampire bats flying near or attacking free-ranging cattle on a pasture.** The behavioral context was derived from synchronized video. We were only able to identify some of the individuals that emitted the calls (when mouth movements were visible on video). Calls include (a) echolocation calls, (b) undulated downsweep, which was only observed in one recording where two bats where flying near a cow, (c) downsweep calls, (d) n-shaped calls, and (e) buzz calls recorded while two bats engaged in antagonistic behavior on a single cow. Inter-call intervals and call sequences have been modified for the figure, except for the call sequence in panel (e), which is a natural sequence recorded from two aggressively interacting bats.

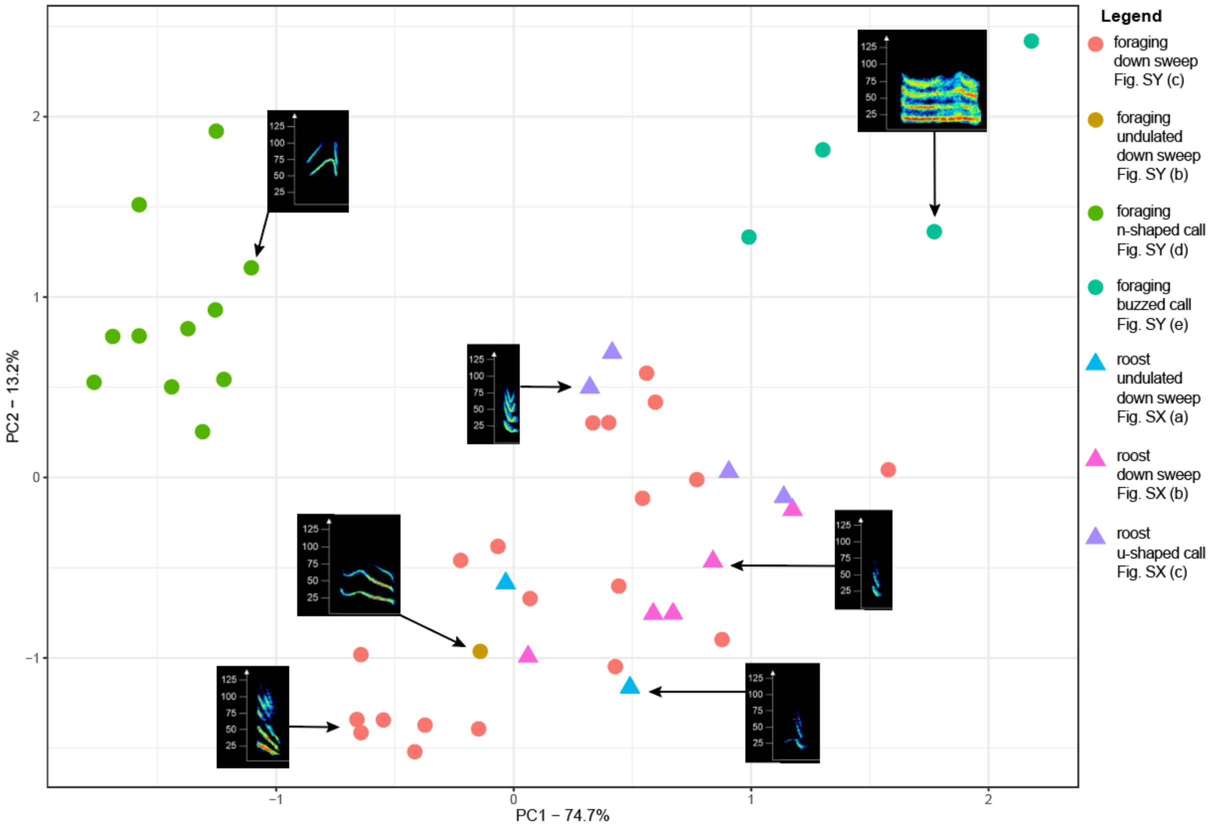


**Figure S7. Variation of call types in multidimensional space.** X and Y axis are first two principal components from spectrum-based parameters measured at 11 positions across each call. Buzz calls produced in antagonistic context and the n-shaped calls were produced during foraging and were distinct from the remaining call types in multivariate space. The legend refers to spectrograms of the calls in Supplementary figures S6 & S7. See Supplementary Table S2 for call parameter measurements.

**Table S1. Within-day effects of roost association rates on foraging encounter rates paired for each daytime period and subsequent night.** Data from day nine was lost due to equipment malfunction. A within-day effect was only detected in day 4, but the model beta coefficients across the 9 days was overall biased above zero (as shown by nonparametric bootstrapping).

| day/night | association | Beta statistic | p | N |
|---|---|---|---|---|
| 2 | 50 cm | -0.001 | 0.98 | 46 |
| | 2cm | 0.016 | 0.57 | 46 |
| 3 | 50 cm | 0.035 | 0.13 | 48 |
| | 2cm | 0.010 | 0.68 | 48 |
| 4 | 50 cm | 0.098 | 0.001* | 46 |
| | 2cm | 0.084 | 0.013* | 45 |
| 5 | 50 cm | 0.047 | 0.11 | 45 |
| | 2cm | 0.001 | 0.96 | 44 |
| 6 | 50 cm | 0.027 | 0.28 | 41 |
| | 2cm | 0.003 | 0.91 | 40 |
| 7 | 50 cm | 0.030 | 0.36 | 39 |
| | 2cm | 0.032 | 0.35 | 39 |
| 8 | 50 cm | -0.021 | 0.33 | 38 |
| | 2cm | -0.009 | 0.66 | 37 |
| 10 | 50 cm | -0.007 | 0.64 | 37 |
| | 2 cm | 0.009 | 0.64 | 34 |
| Mean | 50 cm | 0.026 [0.004 to 0.051] | | |
| [95% CI] | 2 cm | 0.018 [0.003 to 0.04] | | |

**Table S2. Mean social call parameters.** Parameters include call duration, peak frequency of maximum amplitude, minimum fundamental frequency, and maximum fundamental frequency for call types recorded during foraging and inside the roost. The standard deviation is given in parentheses.

| Context | Call type | n | Call duration | Peak frequency | Minimum frequency | Maximum frequency |
|---|---|---|---|---|---|---|
| Foraging | Down sweep | 20 | 4.19 (2.25) | 24.37 (3.93) | 19.64 (3.37) | 33.14 (3.59) |
| | Undulated down sweep | 1 | 20.09 | 23.40 | 17.50 | 34.10 |
| | n-shaped call | 11 | 8.99 (1.42) | 32.87 (2.09) | 27.23 (1.30) | 36.41 (1.33) |
| | Buzz call | 4 | 13.21 (11.68) | 16.80 (3.26) | 13.15 (3.06) | 22.43 (1.79) |
| Roost | Undulated down sweep | 2 | 28.15 (3.61) | 19.00 (0.99) | 17.05 (1.06) | 31.35 (0.49) |
| | u-shaped call | 4 | 20.88 (4.73) | 22.23 (2.36) | 18.00 (2.25) | 32.48 (1.11) |
| | Down sweep | 5 | 11.32 (2.13) | 20.44 (1.82) | 17.38 (2.26) | 32.38 (2.11) |

**Table S3: Summary of behavioral context for social calls produced by vampire bats during foraging.** n = n-shaped call, ds = down sweep, z = buzz call, uds = undulated down sweep; consecutive calls were recorded within less than a second (-) or more than one second (---). Since the recordings were obtained from freely moving animals, some calls were very faint and call sequences may therefore miss individual calls.

| | | | **Number calls per call type** | | | | |
|---|---|---|---|---|---|---|---|
| **Behavioral context bats** | **Behavioral context cows** | **Position of vocalizing bat** | **n** | **ds** | **z** | **uds** | **Call sequence** |
| two bats feeding on one cow | cow moving head, scratching itself | on cow for first two calls then invisible | 0 | 6 | 0 | 0 | ds---ds---ds---ds---ds-ds |
| one bat approaching cow in flight, second bat flies up behind cow; bats circle in flight and one bat lands on cow | grazing calmly | unclear | 0 | 0 | 0 | 2 | uds-uds |
| 1 bat on cow, 2nd bat circles around flying, lands and takes off immediately | grazing calmly | on cow | 3 | 1 | 0 | 0 | n-n-n-ds |
| 1 bat on cow, 1 flying | grazing calmly | unclear | 1 | 0 | 0 | 0 | n |
| 3 bats feeding on 3 cows within few meters (see video S1) | grazing calmly | all three bats on cows vocalize sporadically | 10 | 0 | 0 | 0 | n---n---n-n-n---n---n---n---n---n |
| 3 bats on 3 cows, 1 bat flying by | grazing calmly | Unclear | 1 | 1 | 0 | 0 | ds-n |
| 1 bat on 1 cow | grazing calmly | Unclear | 1 | 0 | 0 | 0 | n |
| 1 bat feeding on cow, second bat flies by, lands, both engage in fight and fly away (see video S2) | grazing calmly | on cow | 3 | 2 | 3 | 0 | n-n-n-ds-z-z-z---ds |
| 1 bat flying | grazing calmly | Unclear | 0 | 3 | 0 | 0 | ds---ds-ds |
| 2 bats feeding on 1 cow | grazing calmly | on cow | 5 | 0 | 0 | 0 | n---n---n-n-n |
| 2 bats feeding on 1 cow | cow moves ear disturbing 1 feeding bat | disturbed bat on cow | 3 | 1 | 0 | 0 | n-ds-n-n |
| 2 bats feeding on 1 cow; 1 bat drinking, second bat moving around, making body contact with first bat, gets hit by the ear of the cow, then both bats start pushing each other from one side of the cow neck to the other side (see video S3) | grazing calmly; slaps a bat with ear | on cow | 5 | 8 | 1 | 0 | n-ds---n-n-n---ds-ds-ds---ds---ds-ds-n-ds-z |
| same 2 bats feeding on same cow (separate wounds) | grazing calmly, walking slowly | on cow | 6 | 2 | 0 | 0 | n-ds-n---n---n-n-n-ds |
| 2 bats on one cow; 1 flies off, returns, finally both fly off (see video S4) | cow is running | bats on cow | 4 | 3 | 0 | 0 | n-n-n---n-n---n-ds-ds-ds |

For the review process, supplementary videos can be accessed under the following link:

https://www.dropbox.com/sh/fuuihw2aohkd0go/AABWTDYsHaEC-o9ZtZ46IjRMa?dl=0

**Video S1:** Three cows are grazing within few meters distance. Each of the three cows has a vampire attached to its neck. Two of the bat individuals seem to be vocalizing in the direction of the other individuals (seconds 1-5 and 23-24).

**Video S2:** A vampire bat seems to be making a bite on the neck of a cow. A second vampire bat joins and both engage in fight and fly away.

**Video S3:** One bat is drinking from an open wound on the neck of a cow. The feet of a second bat hanging on the opposite site of the neck are visible. The first bat moves around, and both bats make body contact. The first bat gets hit by the ear of the cow, then both bats start pushing each other from one side of the cow neck to the other side and a social call is audible (second 28; likely a ‘z’-call).

**Video S4:** Two bats feed from different wounds on the same cow. The cow starts walking towards a second cow. One bat flies up and returns. When the first cow gets pushed by the second cow, the bats fly away.

**Video S5:** Two bats feed from different wounds on the same cow, and one bat vocalizes but not in the direction of the other bat.
