## Supplementary material for "Social foraging in vampire bats is predicted by long-term cooperative relationships": ESM2

Time of foraging bouts by bat and day

Each row shows a bat on a specific date (20–29). Points are encounters within bouts.

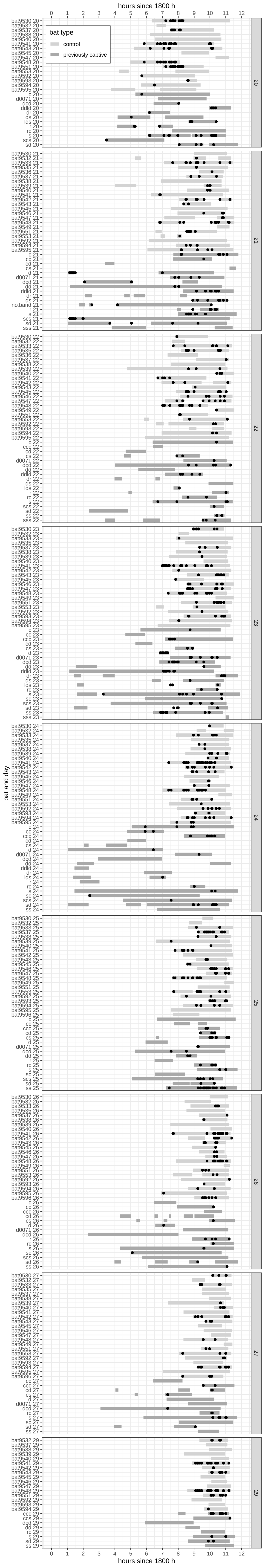
